# *Lactococcus lactis* subsp. *cremoris* C60 promotes immunoglobulin A production from B cells through functional modification of dendritic cells in intestinal environment

**DOI:** 10.1101/2024.10.25.620378

**Authors:** Suguru Saito, Alato Okuno, Nanae Kakizaki, Toshio Maekawa, Noriko M. Tsuji

## Abstract

Probiotics utilizing lactic acid bacteria (LAB) have gained considerable attention in recent trends promoting self-managed health. Specifically, LAB-mediated immune modulation has been a key focus due to its potential to reduce the risk of pathogenic invasion and enhance host immunity. Immunoglobulin A (IgA) plays a crucial role in innate defense against various pathogens. Although only a limited number of probiotic LAB strains have been shown to increase IgA production, this capability remains of great interest. Here, we report a new strain, *Lactococcus lactis* subsp. *cremoris* C60 (C60), which promotes IgA production through functional modulation of intestinal B cells and dendritic cells (DCs). Heat-killed (HK)-C60 increased both pro-inflammatory and anti-inflammatory cytokine productions in DCs via the Toll-like receptor (TLR)-Myeloid differentiation primary response 88 (MyD88) signaling pathway, as demonstrated in both a physiological mouse model and *in vitro* cultures using bone marrow-derived dendritic cells (BMDCs). Notably, intragastric administration of HK-C60 significantly increased systemic IgA production, which was associated with the functional modification of B cells in the Peyer’s patches (PPs) of the small intestine. Immunophenotyping of PP cells from HK-C60-administered mice revealed both an expansion and functional upregulation of B cells. Mechanistically, we identified that DC-derived interleukin-6 (IL-6) and IL-10 play essential roles in the C60-mediated increase in IgA production by B cells. Finally, we evaluated the effect of C60 on IgA production using human peripheral blood mononuclear cells (PBMCs). Consistent with our findings in the mouse model, PBMCs produced IgA upon HK-C60 stimulation in an IL-6- and IL-10-dependent manner. Our results suggest that C60 is a novel probiotic strain capable of promoting intestinal immune homeostasis by upregulating IgA production, underscoring its potential for probiotic applications focused on immune conditioning.

## Introduction

Lactic acid bacteria (LAB) are widely utilized in probiotics to modulate host immune responses [1, 2]. Probiotic LAB primarily modulate immune cell function, conditioning homeostasis and enhancing the immune system’s ability to suppress inflammation and increase resistance to pathogenic invasion [3, 4, 5]. We previously reported that *Lactococcus lactis* subsp. *cremoris* C60 (C60) induces functional modifications in myeloid cells, such as dendritic cells (DCs) and macrophages, resulting in enhanced T cell activity [6, 7, 8]. The probiotic effects of C60 on immunity are based on increased pro-inflammatory cytokine production and enhanced antigen-presenting ability in myeloid cells, both of which promote the differentiation of effector CD4^+^ T cells and strengthen their functions [5, 6]. Moreover, C60 enhances protein antigen processing by activating the 20S immunoproteasome (20SI) and increases major histocompatibility complex (MHC) class I-restricted antigen presentation to CD8^+^ T cells. This modification of DC function leads to increased CD8^+^ T cell anti-tumor activity, resulting in tumor suppression [8].

Although these findings provide sufficient evidence to illustrate C60’s potential for probiotic use, we sought to explore additional immune-related effects of this strain. Myeloid cells often act as key orchestrators in the immune system, and their functional modification by probiotic LAB frequently influences adaptive immunity beyond T cells, impacting other cell types as well. Previous studies have demonstrated that DCs play an important role in B cell activation, specifically in elevating immunoglobulin A (IgA) production [9, 10]. Immunoglobulins (Igs) are key components of humoral immunity and are classified into five types—IgG, IgA, IgM, IgD, and IgE—based on their structure and function [11]. While IgG plays a central role in pathogenic defense due to its superior affinity and specificity, antigen-specific IgG production can take several days following primary infection or antigen recognition, even in individuals previously exposed to the pathogen or vaccinated [12, 13]. During this initial period, IgA provides a crucial first line of defense by working in concert with phagocytic innate immune cells [14]. Although IgA has lower target specificity than IgG, maintaining sufficient IgA levels is critical for increasing the frequency of pathogen capture and elimination [14]. This is especially important in mucosal environments, which are frequently exposed to exogenous factors, including pathogens [15].

While the ability of probiotic LAB to increase IgA production is well-documented, the number of LAB strains with this capability remains limited, highlighting the need for identifying new strains with IgA-enhancing properties.

In this study, we demonstrate that C60 promotes IgA production in B cells by increasing interleukin-6 (IL-6) and IL-10 production in DCs. While we had already established that C60 enhances DC function to bolster CD4^+^ T cell activity, we further investigated whether C60 exerts additional roles in DC immunobiology. We discovered that C60 exhibits a unique characteristic of increasing both pro-inflammatory (Tumor necrosis factor-alpha; TNF-α, IL-6, IL-12) and anti-inflammatory (IL-10) cytokines. Although B cells were directly proliferated by C60’s stimulatory signal, IgA upregulation required IL-6 and IL-10 produced by functionally enhanced DCs. Furthermore, C60 increased IgA production in human peripheral blood mononuclear cells (PBMCs) through the same mechanism identified in the mouse model.

Our findings suggest that C60 is a promising new strain of probiotic LAB capable of elevating IgA levels through immune modulation of DCs, thus contributing to enhanced mucosal immunity.

## Materials and methods

### Reagents and antibodies

Anti-CD45 monoclonal antibody (mAb) (30-F11), anti-CD3ε mAb (17A2), anti-B220 mAb (RA3-6B2), anti-CD138 mAb (281-2), anti-CD11c mAb (N418), anti-IL-6 mAb (MP5-20F3), anti-IL-10 mAb (JES5-16E3), purified anti-CD16/CD32 (2.4G2) mAb, Ultra-LEAF^TM^ purified anti-human IL-6 mAb (MQ2-13A5) Ultra-LEAF^TM^ purified anti-human IL-10 mAb (JES3-9D7) and 7-Aminoactinomycin D (7-AAD) were purchased from BioLegend (San Diego, CA, USA). Anti-IgA mAb (mA-6E1) was purchased from Thermo Fisher Scientific (Waltham, MA, USA). Anti-mouse IL-6 mAb (MP5-20F3), anti-mouse IL-10 mAb (JES5-2A5) were purchased from BioXCell (Lebanon, NH, USA). Corresponding isotype antibodies were purchased from the same companies, respectively. Recombinant murine granulocyte-macrophage colony-stimulating factor (rmGM-CSF) was purchased from Peprotech (Cranbury, NJ, USA). Pam3CysSerLys4 (Pam3CSK4), Pam2CGDPKHPKSF (FSL-1), peptidoglycan (PGN), polyinosinic-polycytidylic acid (Poly (I:C)), Lipopolysaccharide (LPS), imiquimod (IMQ as well as R837), Resiquimod (R848) and ODN 1586 (Class A CpG oligonucleotide) were purchased from Invivogen (San Diego, CA, USA).

### Culture of lactic acid bacteria

*Lactococcus lactis* subsp. *cremoris* C60 (C60) was provided from National Agriculture and Food Research Organization (NARO). C60 was cultured by following the method described in a previous report [6, 7, 8]. *Lactococcus lactis* subsp. *cremoris* SK-11 and HP were both obtained from American Type Culture Collection (ATCC, Manassas, VA, USA) and were cultured by following product data sheet. Bacterial colony forming unit (CFU/mL) was calculated in each culture. For Heat-killed (HK)-LAB preparation, the bacteria were heated at 95°C for 10 min, then the bacterial cells were precipitated by centrifugation at 5,000 *g* for 10 min at 4°C. After being washed with phosphate buffered saline (PBS), the cell pellet was resuspended in saline or PBS and stored at −80°C until use. The sample was used as HK-LAB.

### Animal experiment

C57BL/6 wild-type (WT) mice were purchased from CLEA Japan, Inc. (Tokyo, Japan). Myeloid differentiation primary response 88-knockout (MyD88-KO) mice were obtained from Jackson laboratory (Ibaraki, Japan). All mice were bread under specific pathogen-free conditions with 12 h day/night cycles. Gender-matched 8 to12 weeks old mice were used for experiments. Some mice received intragastric (*i.g.*) administration of saline (200 μL) or HK-C60 (200 μL of 5.0×10^9^ CFU/mL suspension) at every 24 h for 14 days. All experimental protocols were reviewed and approved by the animal welfare committee of National Institute of Advanced Industrial Science and Technology (AIST) (protocol No. 109) and Shibata Gakuen University (protocol No. 2107)

### Preparation of PP-derived cells

PPs were collected from small intestine and the cells were isolated by following a protocol described in previous report [6, 7]. Briefly, the PPs were incubated in epithelial dissociation buffer (PBS containing 10% Fetal Bovine Serum (FBS), 100 mg/mL penicillin, 100 mg/mL streptomycin, 20 mM EDTA and 10 mM dithiothreitol (DTT)) at 37°C for 20 min with staring. Then, the PPs were treated with digestion buffer (RPMI1640 supplemented with 10% FBS, 100 mg/mL penicillin, 100 mg/mL streptomycin, 1 mg/mL collagenase type-I, 50 μg/mL DNase) at 37 °C for 30 min with staring. After digestion, the PPs were collected and crushed on a 70 μm cell strainer. The isolated cells were washed with cell culture medium (RPMI 1640 supplemented with 10% FBS, 50 μM 2-mercaptoethanol (2-ME), 10 mM 4-(2-hydroxyethyl)-1-piperazineethanesulfonic acid (HEPES), 100 mg/mL penicillin, 100 mg/mL streptomycin), then the sample was filtered through a 40 μm cell strainer again to make single cell suspension. After being washed with cell culture medium, the cells were finally collected by centrifugation at 300 *g* for 5 min. The precipitated cells were used as PP-derived cells. The cell viability was assessed by trypan blue staining in each sample and the samples with more than 95% of live cells were used for experiments.

### Preparation of bone marrow derived dendritic cells (BMDCs)

BM cells were isolated from the tibia and femur of naïve WT mice (8-10 weeks) and seeded at 3.0×10^5^/mL in cell culture medium containing rmGM-CSF (20 ng/mL). The cells were cultured at 37°C for 8 days with medium replaced half-volume on days 3 and 6. On day 8, floating cells were harvested as BMDCs and washed with cell culture medium. The purity of differentiated BMDC was assessed by flow cytometry, and the samples with more than 90% of CD11c+ cells were used for subsequent experiment.

### Stimulation of BMDCs for cytokine production assay

BMDC (1.0×10^6^/mL) were seeded in 96 well flat bottom plate with cell culture medium containing vehicle (PBS), Pam3CSK4 (1 ug/mL), PGN (10 ug/mL), FSL-1 (1 ug/mL), Poly (I:C) (25 ug/mL), LPS (1 ug/mL), R837 (25 ug/mL), R848 (10 ug/mL), ODN1585 (10 ug/mL) or HK-C60 (1.0×10^8^, 2.5×10^8^ or 5.0×10^8^ CFU/mL) at 37°C for 24 h. The cultured medium was collected and stored at −80°C until use. The cytokine concentrations in cultured medium were measured by ELISA.

### Stimulation of PP-derived cells and measurement of IgA production

The PP-derived cells (1.0×10^7^/mL) were seeded in 96 well round bottom plate with cell culture medium. The cells were treated with vehicle (PBS), LPS (1 μg/mL) or HK-C60 (1.0×10^8^, 2.5×10^8^ or 5.0×10^8^ CFU/mL) at 37 °C for 72 h. In cytokine neutralizing assay, the cells were treated with anti-mouse IL-6 mAb (10 μg/mL) and/or anti-mouse IL-10 mAb (10 μg/mL). Corresponding isotype antibody was used for control treatment. The cultured medium was collected and stored at −80°C until use. The IgA and cytokine concentrations in cultured medium were measured by ELISA.

### Peripheral blood mononuclear cell (PBMC) culture and stimulation assay

Human PBMCs were obtained from Zen-Bio (Durham, NC, USA). The frozen cell stocks were thawed on ice, then cultured in pre-warmed medium provided by the company. The cells were pre-incubated at 37°C for overnight, then used for stimulation assay. The PBMCs were cultured at 1.0×10^7^/mL in cell culture medium supplemented with vehicle, anti-human IgM (10 μg/mL)+CD40L (1 μg/mL), anti-human IgM (10 μg/mL)+ODN2006 (5 μg/mL), HK-C60 (1.0×10^8^, 2.5×10^8^ or 5.0×10^8^ CFU/mL) at 37°C for 72 h. In cytokine neutralizing assay, the cells were treated with anti-moure IL-6 mAb (10 μg/mL) and/or anti-mouse IL-10 mAb (10 μg/mL). Corresponding isotype antibody was used for control treatment. The IgA concentration in cultured medium was measured by ELISA.

### Reverse Transcription quantitative polymerase chain reaction (RT-qPCR)

The small intestinal PPs were extracted from the mice received saline or HK-C60 by i.g. administration for 14 days. CD11c+ DCs were isolated from whole PP-derived cells using CD11c MicroBeads UltraPure, mouse (Miltenyi Biotec). Total RNA was isolated from the DCs using TRIzol (Thermo Fisher Scientific), then concentration and purity were measured by NanoDrop 2000c (Thermo Fisher Scientific). Total RNA (200 ng) was used for reverse transcription for making complementary DNA (cDNA) using PrimeScript RT Master Mix (Takara, Tokyo, Japan). The cDNA was subjected to quantitative PCR performed by using TB Green Premix Ex Taq (Takara) in Thermal Cycler Dice Real Time System (Takara). The *Gapdh* mRNA expression was used as an internal control. The mRNA expression was quantified by ΔCt method. The primer sequences used in analysis were shown in Supplementary Table 1.

### Flow cytometry

Flow cytometry analysis was performed by using a flow cytometer (Aria I or LSR-II; BD Biosciences, Franklin Lakes, NJ, USA) with the fluorochrome-conjugated monoclonal antibodies (mAbs) described in reagents and antibodies. For surface marker staining, the cells were incubated with FcR blocker (anti-CD16/32; 2.4G2) at 4°C for 10 min followed by incubation with fluorochrome-conjugated mAb at 4°C for 30 min. Intracellular cytokine staining was performed by using BD Cytofix/Cytoperm kit with GolgiStop^TM^. Briefly, extracellular marker-stained samples were fixed and permeabilized at 4°C for 20 min, then intracellular cytokine was stained with fluorochrome-conjugated mAb at 4°C for 30 min. The dead cells were excluded by 7-actinomycin D (7-AAD) staining during analysis. All data were analyzed by BD FACS Diva (BD Bioscience,) or FlowJo (BD Biosciences).

### Enzyme-Linked Immunosorbent Assay (ELISA)

Cytokines concentrations were measured by ELISA using DuoSet ELISA kit (R&D systems) for each target. All procedures were performed by following products manuals.

### Statistical Analyses

GraphPad Prism (GraphPad Software, La Jolla, CA, USA) was used for statistical analysis. The student’s t-test or one-way analysis of variance (ANOVA) were used for comparisons between two groups and multiple groups, respectively. Values of *p* < 0.05, *p* < 0.01, and *p* < 0.001 were considered as statistically significant.

## Results

### C60 Increases production of both pro-inflammatory and anti-inflammatory cytokines in intestinal DCs

While our previous reports demonstrated that C60 induces pro-inflammatory cytokine production in myeloid cells, the range of cytokines examined was limited [6, 7]. To gain a deeper understanding of the cytokine specificity induced by C60, we analyzed the cytokine gene expression profile in dendritic cells (DCs) isolated from the small intestinal PPs of saline-or heat-killed (HK) C60-administered mice. Mice received intragastric (i.g.) administration of either saline or HK-C60 for 14 days, after which CD11c^+^ DCs were isolated from PPs using magnetic separation (Figure 1A). We then assessed the mRNA expression of 13 different cytokines in these DCs using real-time PCR.

**Figure 1.**
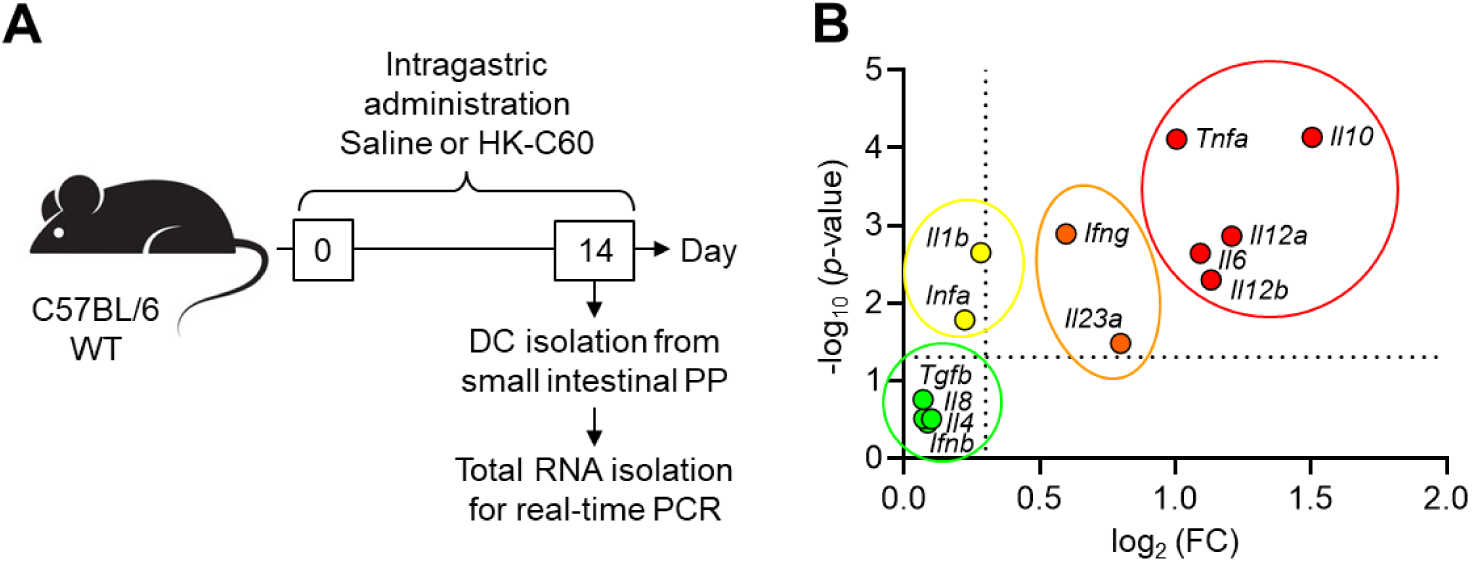
Cytokine gene expression profile in intestinal DCs. A) The experimental design of intragastric (i.g.) administration and cytokine gene expression profiling. The WT mice received i.g. administration of saline or HK-C60 every day for 14 days (n=3 in each group). CD11c^+^ DCs were isolated from small intestinal PPs by using magnetic beads. The total RNA was isolated from DCs and subjected to real-time PCR. B) Cytokine gene expressions in intestinal DCs. The gene expressions were classified to four clusters (green, yellow, orange and red) following the p-value and fold change (FC) calculated by 2^-ΔΔCt^ method. Student t-test was used to analyze data for significant differences. Values of *p* < 0.05 was regarded as significant.

Interestingly, we identified four distinct clusters of cytokine gene expression patterns in response to HK-C60 (Figure 1B). Cluster 1, marked by a green circle, included *Il4*, *Il8*, *Ifnb,* and *Tgfb*, which showed slight but non-significant increases in mRNA expression following HK-C60 stimulation. Cluster 2, marked by a yellow circle, contained *Ifna* and *Il1b*. These cytokines were significantly upregulated in response to HK-C60, though the increase was modest (less than two-fold compared to the control). Clusters 3 and 4, marked by orange and red circles, respectively, contained a total of seven genes: *Ifng*, *Il23A*, *Il6*, *Il12A*, *Il12B*, *Il10*, and *Tnfa*. These cytokines showed significant upregulation, with increases of more than four-fold in HK-C60-treated DCs compared to controls. Notably, the genes in cluster 4 exhibited a particularly strong response, with expression levels increasing more than ten-fold. Among the cytokines, *Il10* showed the highest level of upregulation, with more than a 32-fold increase in mRNA expression. This finding highlights C60’s ability to induce both pro-inflammatory and anti-inflammatory cytokine production in DCs.

### Characterization of cytokine production in HK-C60-stimulated BMDCs

To further explore cytokine production at the protein level, we conducted an in vitro assay using BMDCs. Based on the significant mRNA expression of TNF-α, IL-6, IL-12, and IL-10 observed in intestinal DCs from C60-administered mice, we focused on measuring these cytokines in BMDCs. BMDCs were generated by differentiating bone marrow leukocytes in the presence of GM-CSF for 8 days, followed by stimulation with HK-C60. To assess the relative efficacy of HK-C60 in inducing cytokine production, we included specific Toll-like receptor (TLR) ligands as positive control stimuli in the assay (Figure 2A).

**Figure 2.**
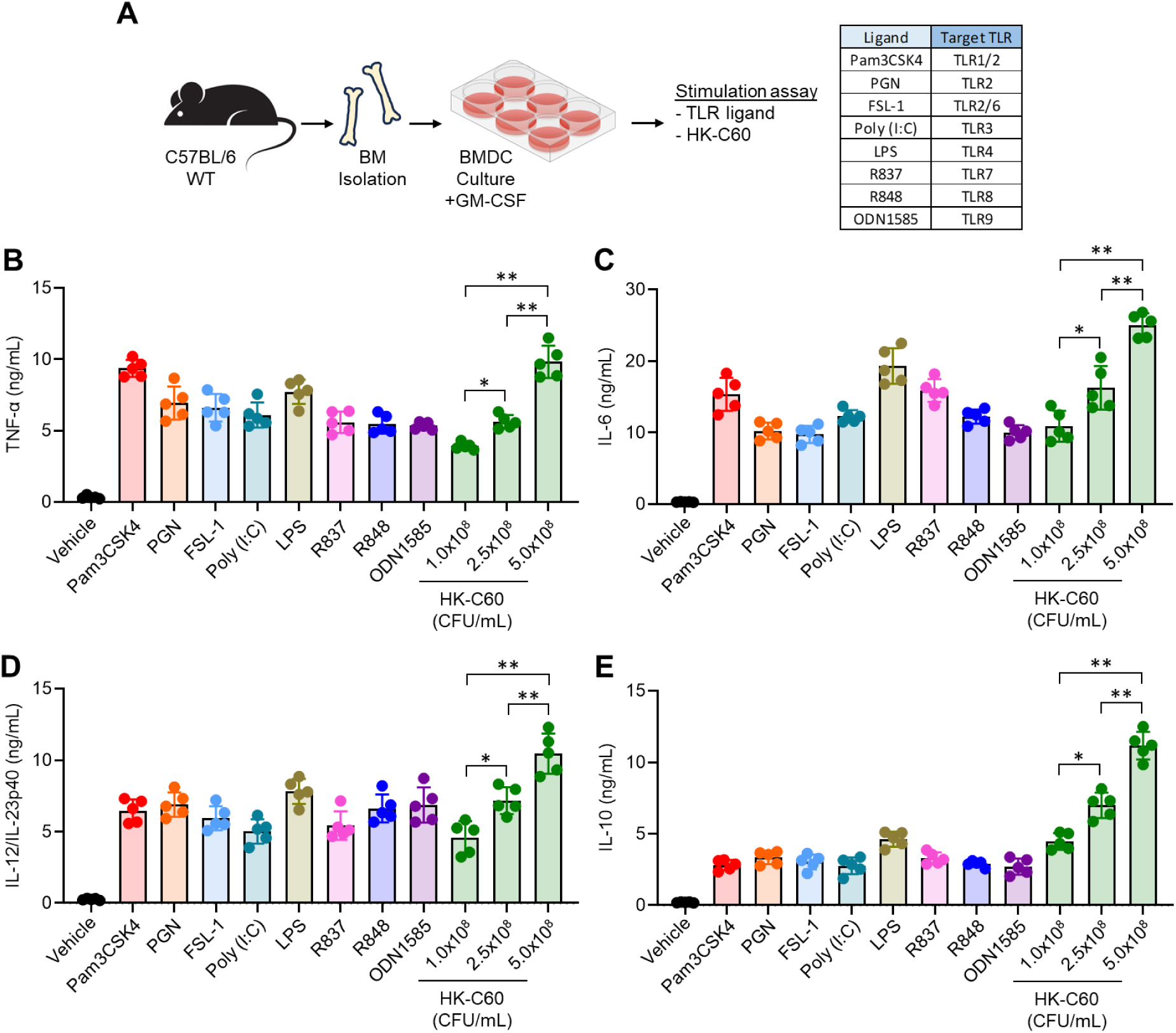
Cytokine production assay in HK-C60-stimulated BMDCs. A) Experimental design of BMDC stimulation assay with HK-C60. BMDCs were prepared by culturing with rmGM-CSF for 8 days (n=5 for independent culture). The differentiated BMDCs were subjected to in vitro stimulation assay with TLR ligand or HK-C60. Used TLR ligands and target receptors are shown in table. B-E) Cytokine production in BMDC stimulation assay. The BMDCs were cultured with vehicle, TLR ligand or HK-C60 at 37C for 24 h. The concentration of TNF-α (B), IL-6 (C), IL-12/IL-23p40 (D) and IL-10 in the conditioned medium were measured by ELISA, respectively. The data were shown as mean ± SEM of five samples from independent experiments. Student t-test or one-way ANOVA was used to analyze data for significant differences. Values of \**p* < 0.05, \*\**p* < 0.01 and \*\*\**p* < 0.001 were regarded as significant. ns: not significant.

All TLR ligands used in this assay elicited 5- to 10-fold higher concentrations of pro-inflammatory cytokines (TNF-α, IL-6, and IL-12/IL-23p40) in BMDCs compared to vehicle controls. Similarly, HK-C60-stimulated BMDCs produced significantly higher levels of these pro-inflammatory cytokines in a dose-dependent manner (Figure 2B-D). IL-10, an anti-inflammatory cytokine, was also significantly elevated in BMDCs stimulated with TLR ligands compared to controls, although its production was lower than that of the pro-inflammatory cytokines. Interestingly, IL-10 production in BMDCs was notably higher in response to HK-C60 stimulation than that of pro-inflammatory cytokines (Figure 2E). Although we had previously compared the pro-inflammatory cytokine production between C60 and other LAB strains in BMDC stimulation assays, IL-10 production had not yet been examined. Similar to the pro-inflammatory response, HK-C60 significantly increased IL-10 levels compared to vehicle controls, whereas the control LAB strains did not induce substantial IL-10 production (Supplemental Figure 1). These findings indicate that C60 is a unique probiotic strain capable of inducing both pro-inflammatory and anti-inflammatory cytokine production in DCs.

### C60 stimulation activates the TLR-MyD88 pathway in cytokine production by DCs

To investigate the mechanism by which C60 induces cytokine production in dendritic cells (DCs), we examined the involvement of the Toll-like receptor (TLR)-MyD88 signaling pathway. Given that lactic acid bacteria (LAB) and their byproducts are often recognized by pattern recognition receptors (PRRs) similar to those used to detect pathogenic bacteria, we assessed the expression of various TLRs and MyD88 in PP-derived DCs obtained in the experiment presented in Figure 1. HK-C60 administration significantly upregulated the mRNA expression of *Tlr2*, *Tlr4*, *Tlr6*, and *Tlr9* in intestinal DCs compared to controls. Additionally, MyD88, an essential downstream adaptor protein for all TLR signaling pathways except for TLR3, was also significantly upregulated in HK-C60-stimulated DCs (Figure 3A).

**Figure 3.**
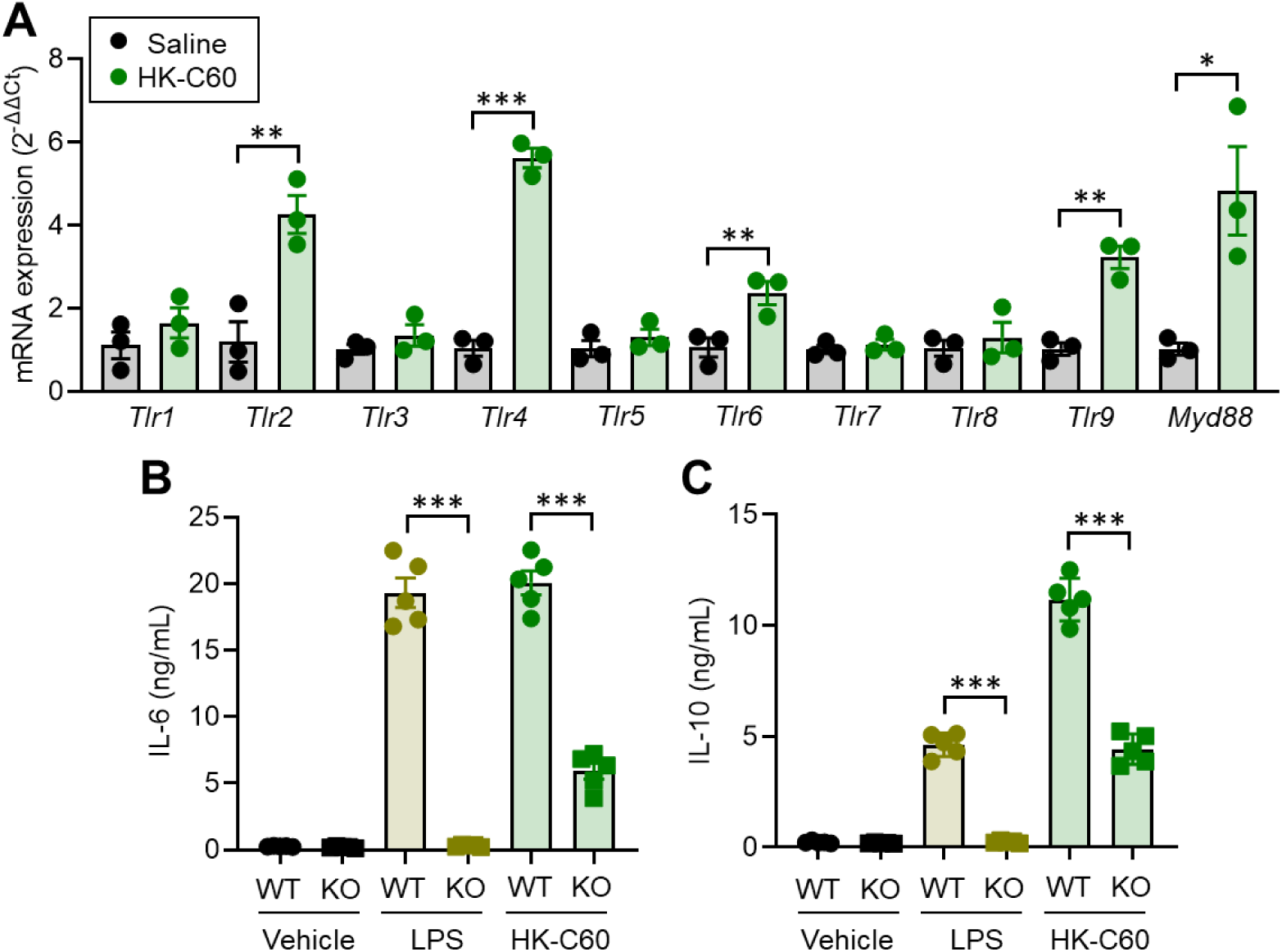
HK-C60 activates cytokine production in DCs via TLR-MyD88 pathway. A) Gene expression profile in PP-derived DCs. The WT mice received saline of HK-C60 i.g. administration for 14 days following the protocol presented in Figure 1A (n=3 in each group). The CD11c^+^ DCs were isolated from small intestinal PP and total RNA was subjected to real-time PCR investigating TLR and MyD88 mRNA expressions. The mRNA expression was quantified by 2^-ΔΔCt^ method. B-C) In vitro BMDC stimulation and cytokine production assay. The BMDCs were prepared from WT or MyD88-KO mice (n=5 in each). The BMDCs were cultured with vehicle, LPS (1 ug/mL) or HK-C60 (5.0×10^8^ CFU/mL) at 37C for 24 h. The concentration of IL-6 (B) and IL-10 (C) in the conditioned medium were measured by ELISA, respectively. The data were shown as mean ± SEM of three to five samples from independent experiments. Student t-test or one-way ANOVA was used to analyze data for significant differences. Values of \**p* < 0.05, \*\**p* < 0.01 and \*\*\**p* < 0.001 were regarded as significant. ns: not significant.

To further investigate the requirement of the TLR-MyD88 pathway in cytokine production induced by HK-C60, we performed in vitro stimulation assays using BMDCs from wild-type (WT) and MyD88 knockout (MyD88-KO) mice. As expected, IL-6 and IL-10 production induced by LPS stimulation in WT BMDCs was completely abolished in MyD88-KO BMDCs (Figure 3B, C, gold). In HK-C60-stimulated BMDCs, the absence of MyD88 did not completely abolish IL-6 and IL-10 production but led to a 60-70% reduction in cytokine levels compared to WT BMDCs (Figure 3B, C). These results indicate that the TLR-MyD88 signaling pathway plays a predominant role in cytokine production induced by HK-C60 stimulation in DCs.

### HK-C60 administration increases IgA production in mice

Probiotic LAB are expected to condition immune responses, and we aimed to explore additional functions of C60 beyond the previously reported T cell functional modulation [6, 7, 8]. To investigate the immunological effects of C60 in the intestinal environment, we administered saline or HK-C60 by intragastrically (i.g.) to mice for 14 days, following the schedule outlined in Figure 1A. After the treatment, cells from the small intestinal PPs were isolated and analyzed using flow cytometry. A global characterization of immune cells using t-distributed stochastic neighbor embedding (t-SNE) plots revealed a significant increase in the B220^+^ B cell population in HK-C60-treated mice compared to controls, suggesting notable changes in B cell activity. In contrast, no significant populational changes were observed in other cell markers, such as CD3^+^ (T cells), CD11c^+^ (dendritic cells), CD11b^+^ (common myeloid cells), F4/80^+^ (macrophages), Ly-6C^+^ (monocytes), or Ly-6G^+^ (neutrophils) (Figure 4A). Further analysis of cell populations in t-SNE plots highlighted three clusters (red, blue, and orange) that were predominantly expanded in HK-C60-treated mice. These clusters consisted mainly of B220^+^ cells, indicating a major shift in B cell populations (Supplemental Figure 2 A, B). Upon closer examination, B cells in the PPs of HK-C60-treated mice exhibited an activated status, as demonstrated by significantly increased expression of activation markers CD80, CD86, and major histocompatibility complex (MHC) class II (Figure 4B-E).

**Figure 4.**
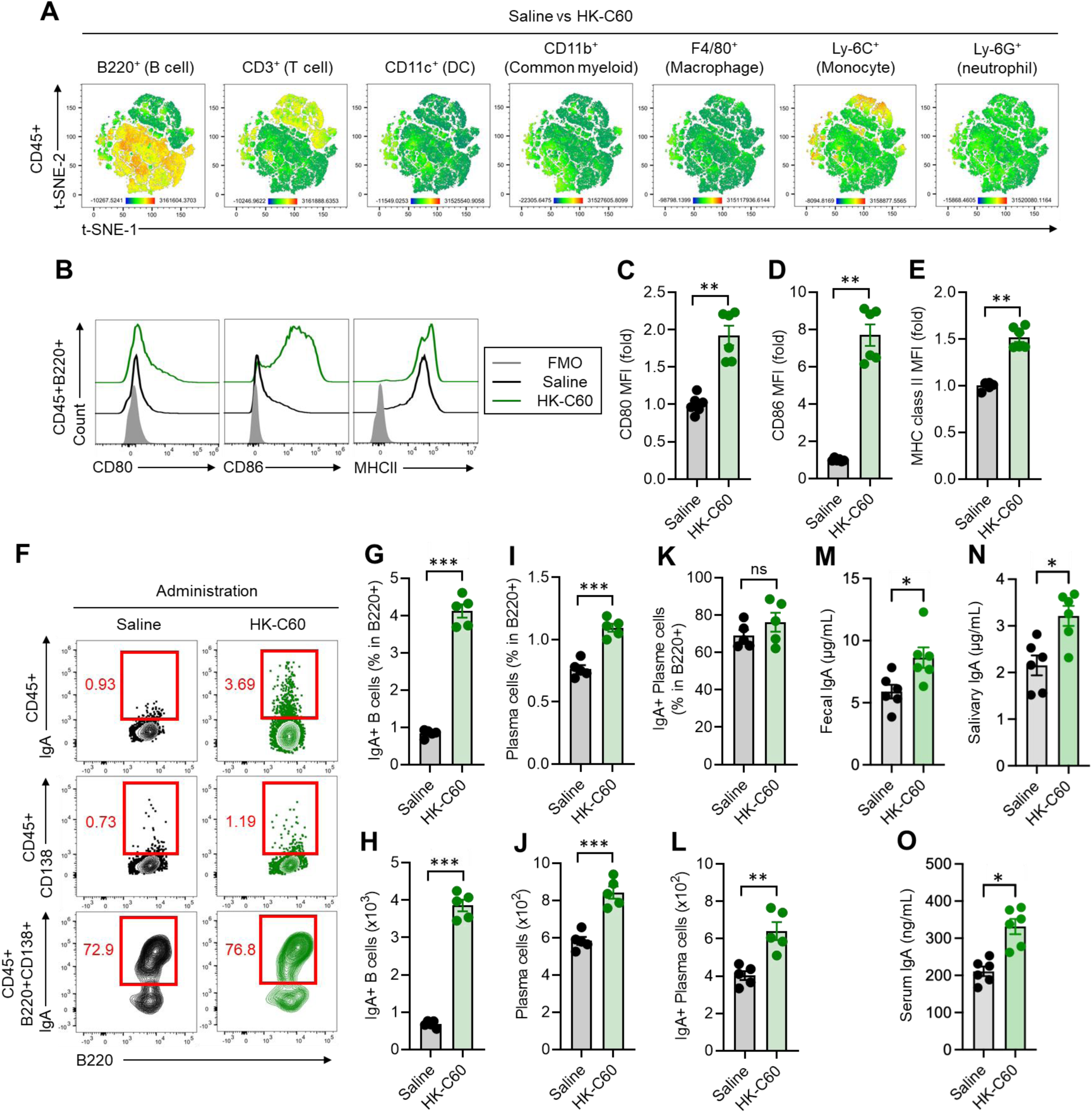
HK-C60 administration increases IgA production from intestinal B cells. The WT mice received saline of HK-C60 i.g. administration for 14 days following the protocol presented in Figure 1A (n=6 in each group). The small intestinal PP cells were isolated and subjected to flow cytometry analysis. Feces, saliva and plasma samples were also correct from the mice is day 14 of treatment and used for IgA ELSIA assay. A) The difference between saline and HK-C60-administered mice in each immune cell type presented by t-SNE. B-C) B cell functional analysis. B) Representative histograms of CD80, CD86 and MHC class II and Cumulative MFI values (fold change to control) of CD80 (C), CD86 (D) and MHC class (E) in B220^+^ B cells. F-L) IgA expression in B cells. F) Representative plots of IgA expression in total B220^+^ B cells (top), CD138^+^ plasma cells (middle) and IgA expression of CD138^+^ plasma cells (bottom). G) Percentage of IgA^+^B220^+^ B cells. H) Cell number of IgA^+^B220^+^ B cells. I) Percentage of CD138^+^ plasma cells. J) Cell number of CD138^+^ plasma cells. K) Percentage of IgA+CD138^+^ plasma cells. L) Cell number of IgA^+^CD138^+^ plasma cells. The cell numbers are described as per 10^5^ of CD45^+^ cells. The data were shown as mean ± SEM of three to five samples from independent experiments. Student t-test or one-way ANOVA was used to analyze data for significant differences. Values of \**p* < 0.05, \*\**p* < 0.01 and \*\*\**p* < 0.001 were regarded as significant. ns: not significant.

Next, we assessed B cell function and found that IgA production was significantly enhanced in HK-C60-administered mice. The frequency of IgA-expressing B cells in PPs was markedly higher in the HK-C60 group (4.120 ± 0.398%) compared to controls (0.838 ± 0.101%) (Figure 4F top, 4G). Additionally, the total number of IgA^+^ B cells was significantly elevated (saline: 0.679 ± 0.086 × 10³ vs. HK-C60: 3.854 ± 0.347 × 10³ per 10⁵ CD45^+^ cells) (Figure 4H). Plasma cell (PC) numbers were also increased in HK-C60-treated mice. The percentage of CD138^+^ plasma cells within the B cell population rose significantly (saline: 0.764 ± 0.067% vs. HK-C60: 1.096 ± 0.070%) (Figure 4F mid, 4I), and the number of PCs increased from 5.823 ± 0.474 × 10² in saline-treated mice to 8.417 ± 0.737 × 10² per 10⁵ CD45^+^ cells in HK-C60-administered mice (Figure 4J). While the percentage of IgA^+^ plasma cells showed a slight increase in HK-C60-treated mice, the difference was not statistically significant (saline: 69.06 ± 6.674% vs. HK-C60: 76.16 ± 11.43%) (Figure 4F low, 4K). However, the number of IgA^+^ plasma cells was significantly elevated in the HK-C60 group (saline: 4.028 ± 0.527 × 10² vs. HK-C60: 6.408 ± 1.078 × 10² per 10⁵ CD45+ cells) (Figure 4L).

The upregulation of IgA by HK-C60 was not confined to the intestinal environment but also observed systemically. IgA concentrations in feces were significantly higher in HK-C60-treated mice compared to controls (Figure 4M). Additionally, both saliva and plasma samples from HK-C60-administered mice exhibited increased IgA levels compared to saline-treated mice (Figure 4N, O).

We also investigated whether the abolition of TLR signaling affects IgA production in PP-derived cells. The PP-derived cells from naive MyD88-KO mice exhibited significantly decreased IgA production compared to equivalent WT cells upon LPS and HK-C60 stimulation (Supplemental Figure 3). These findings demonstrate that HK-C60 enhances IgA production from intestinal B cells, promoting both local and systemic immune defenses.

### DC-derived IL-6 and IL-10 play crucial roles in HK-C60-induced IgA upregulation

To investigate the mechanism behind HK-C60-induced IgA upregulation, we established an in vitro culture system of PP)-derived cells to assess whether HK-C60 can stimulate IgA production outside of physiological conditions. We conducted two different experiments. In the first experiment, PP-derived cells were isolated from naïve WT mice and cultured with vehicle (PBS), LPS, or HK-C60 at 37°C for 72 hours (Figure 5A, Exp 1). In the second experiment, WT mice received intragastric (i.g.) administration of saline or HK-C60, and then PP-derived cells were cultured under the same conditions as in Exp 1 (Figure 5A, Exp 2). IgA concentrations in the cultures were measured using ELISA. In Exp 1, LPS stimulation induced IgA production, and HK-C60 further increased IgA production in a dose-dependent manner, peaking at 5.0 × 10⁸ CFU/mL (MOI = 1:50) (Figure 5B). The pre-administration of HK-C60 in Exp 2 enhanced IgA production in PP-derived cultures, with both LPS and HK-C60 stimulating greater IgA production compared to Exp 1 (Figure 5C). Notably, other Lactococcus lactis subsp. cremoris strains, SK-11 and HP, did not significantly induce IgA production in PP-derived cultures from naïve WT mice (Supplemental Figure 4), suggesting that C60 is a unique probiotic strain capable of efficiently inducing IgA production in the intestinal environment.

**Figure 5.**
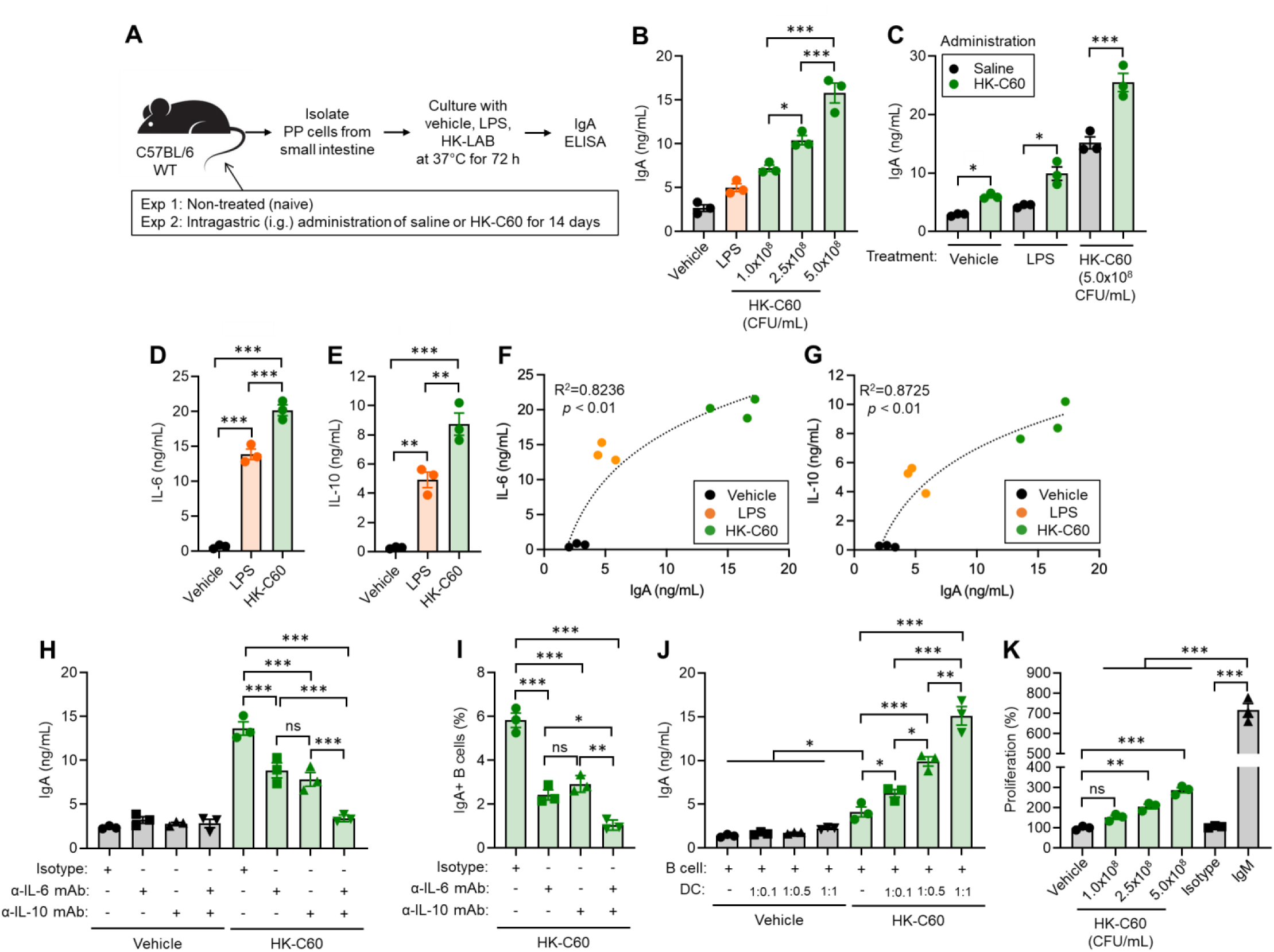
HK-C60-derived IgA upregulation is DC cytokine dependent manner. A) The experimental design of IgA production assay using PP-derived cells. The PP-derived cells were prepared from two different experimental conditions (Exp 1 and Exp 2). In Exp 1, The cells were isolated from small intestinal PPs of naive C57BL/6 WT mice (n=3). In Exp 2, the PP cells were isolated from saline (200 μl) or HK-C60 (200 μl of 5.0×10^9^ CFU/mL) administered C57BL/6 WT mice (n=3 in each group). The isolated cells (1.0×10^7^/mL) were cultured with vehicle (PBS), LPS (1 ug/mL, positive control) or HK-C60 (indicated does) at 37°C for 72 h, then IgA concentration in the cultured medium was measured by ELISA. IgA concentrations were represented in B) Exp 1 and C) Exp 2, respectively. D-E) Cytokine productions in PP-derived cell cultures. The cultures were prepared by WT naïve mice (n=3) following Exp 1 condition, and treated with vehicle (PBS), LPS (1 ug/mL) or HK-C60 (5.0×10^8^ CFU/mL) at 37C for 72 h. The IL-6 (D) and IL-10 (E) concentrations in cultured medium were measured by ELISA, respectively. F-G) Correlation of IL-6 and IgA concentration (F) and IL-10 and IgA (G) in the cultured medium obtained from culture condition (D) and (E). H) In vitro cytokine neutralization assay. The PP-derived cells were prepared from WT mice (n=3) and the cells (1.0×10^7^/mL) were treated with vehicle (PBS) or HK-C60 (5.0×10^8^ CFU/mL). The culture was further treated with anti-IL-6 and/or IL-10 mAb (both 10 μg/mL; +) or isotype antibody (10 μg/mL; −) at 37°C for 72 h. The IgA concentration in the cultured medium was measured by ELISA. I) The percentage of IgA+B220+ B cells in the culture established in (H) (HK-C60-treated group only). J) IgA production assay in B cell and DC coculture. Both B cells and DCs were isolated form small intestinal PPs of naïve WT mice. The B cells (1.0×10^7^/mL) were seeded in the plate alone or together with DC at indicated rations (B cell:DC=1:0.1, 1:0.5, or 1:1). The cultures were further treated with vehicle (PBS) or HK-C60 (5.0×10^7^ CFU/mL for 1:0.1, 2.5×10^7^ CFU/mL for 1;0.5 or 5.0×10^8^ CFU/mL for 1:1). The cultures were incubated at 37C for 72 h, then IgA concentrations in cultured medium were measured by ELISA. K) B cell proliferation assay. PP-derived B cells (1.0×10^7^/mL) were cultured with vehicle (PBS), HK-C60 (indicated doses), isotype antibody (10 ug/mL) or anti-IgM mAb (10 ug/mL) at 37C for 72 h. The B cell proliferation was measured by MTT assay. The data were shown as mean ± SEM of three samples. One-way ANOVA was used to analyze data for significant differences. Values of \**p* < 0.05, \*\**p* < 0.01 and \*\*\**p* < 0.001 were regarded as significant. ns: not significant.

We then explored the role of DCs in promoting IgA production through cytokine modulation. Our previous studies showed that LAB-induced IL-6 and IL-10 are essential for T cell-independent IgA production, with both cytokines produced by DCs rather than B cells [8]. To investigate whether IL-6 and IL-10 are also crucial in the HK-C60-stimulated PP-derived culture (Exp 1, Figure 5A), we measured their concentrations. Both cytokines were significantly elevated in HK-C60-stimulated cultures (Figure 5D, E), and their levels correlated positively with IgA production (Figure 5F, G).

To confirm the crucial roles of IL-6 and IL-10 in IgA upregulation, we performed cytokine neutralization using monoclonal antibodies (mAbs) in PP-derived cultures with HK-C60 stimulation. Neutralization of either IL-6 or IL-10 significantly reduced IgA production compared to isotype antibody controls, while neutralization of both cytokines further reduced IgA levels to those observed in isotype antibody-treated cultures (Figure 5H). Similarly, IL-6 or IL-10 neutralization decreased the frequency of IgA^+^ B cells in HK-C60-stimulated cultures (Figure 5I).

To confirm the indispensable role of DCs in cytokine-dependent IgA production, we tested IgA production in different culture conditions: B cells alone and B cells co-cultured with DCs. Both cell types were isolated from small intestinal PPs of naïve WT mice. B cells cultured alone produced slightly more IgA than controls upon HK-C60 stimulation, but the increase was not statistically significant. However, in co-culture with DCs, IgA production from B cells significantly increased upon HK-C60 stimulation. Without HK-C60 stimulation, DCs did not induce IgA production from B cells (Figure 5J). These results demonstrate that DC-derived IL-6 and IL-10 play indispensable roles in IgA production from B cells in response to HK-C60 stimulation. Although B cell alone did not produce significant level of IgA, HK-C60 stimulation significantly increased B cell proliferation, suggesting that this direct expansion of the B cell population may also contribute to the upregulation of IgA production (Figure 5K).

### C60 increases IgA production in human PBMCs

To investigate whether C60 has a similar effect on IgA upregulation in humans as observed in mice, we performed an in vitro stimulation assay using human peripheral blood mononuclear cells (PBMCs). The positive controls (anti-IgM + CD40L and anti-IgM + ODN2006) significantly increased IgA production in the cultures compared to the vehicle-treated samples. Notably, HK-C60 stimulation further enhanced IgA production in a dose-dependent manner, mirroring the effects observed in murine PP-derived cultures (Figure 6A). We also conducted a cytokine neutralization assay in human PBMC cultures. Neutralization of IL-6 or IL-10 independently decreased IgA production in HK-C60-stimulated cultures; however, IgA levels remained significantly higher compared to vehicle treatment. In contrast, neutralization of both IL-6 and IL-10 abolished IgA production induced by HK-C60 stimulation in human PBMCs (Figure 6B). Thus, the effect of HK-C60 on IgA induction is conserved between mice and humans.

**Figure 6.**
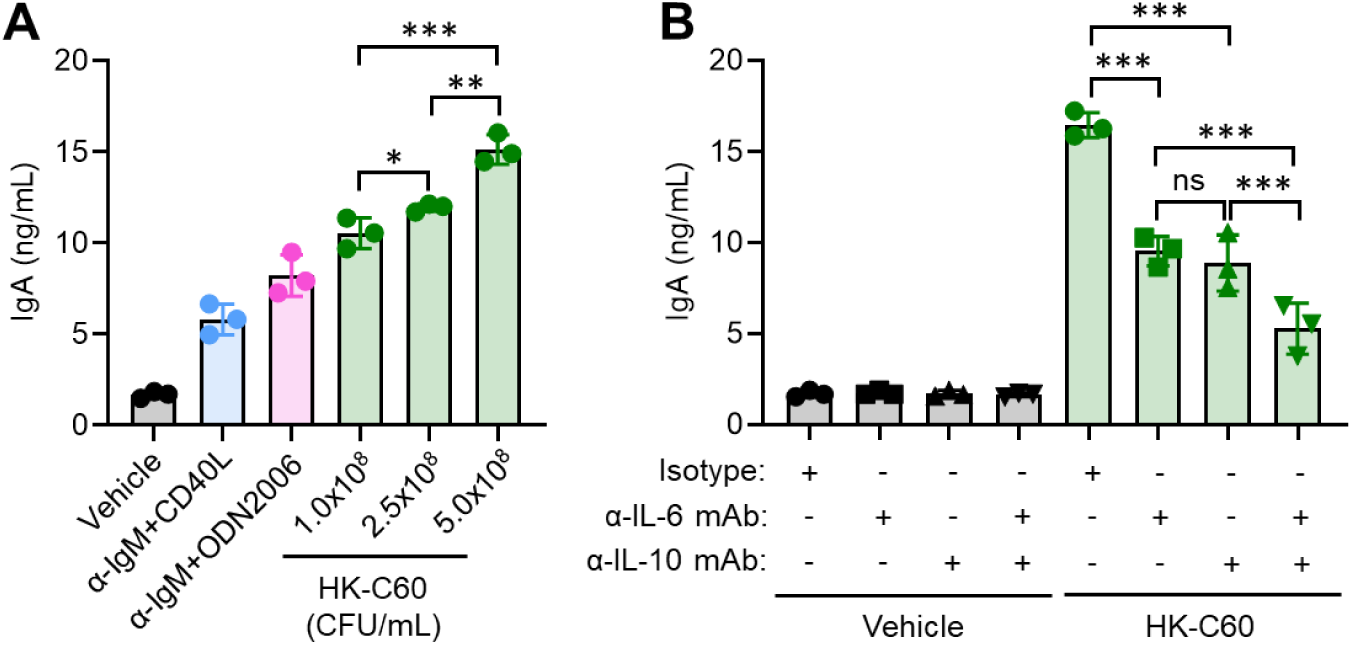
HK-C60 produces IgA in human PBMCs. G) IgA production in human PBMCs. PBMCs (1.0×107/mL) were cultured with vehicle (PBS), anti-IgM (10 μg/mL)+CD40L (1 μg/mL), anti-IgM (10 μg/mL)+ODN2006 (5 μg/mL), HK-C60 (1.0×108, 2.5×108 or 5.0×108 CFU/mL) at 37°C for 72 h. The IgA concentration in cultured medium was measured by ELISA. The data were shown as mean ± standard error mean (SEM) of three or six samples. Student t-test or one-way ANOVA was used to analyze data for significant differences. Values of \**p* < 0.05, \*\**p* < 0.01 and \*\*\**p* < 0.001 were regarded as significant. ns: not significant.

## Discussion

This study demonstrated that C60 is capable of inducing a significant production of both pro-inflammatory and anti-inflammatory cytokines from DCs. Specifically, the levels of TNF-α, IL-6, IL-12, and IL-10 were markedly elevated in DCs in response to bacterial stimuli. This is a unique characteristic of the C60 strain, distinguishing it from other probiotic LAB strains. While the upregulation of pro-inflammatory cytokines is a common phenomenon in the immune response to probiotic LAB, the significant increase in IL-10 is less frequent. IL-10 plays a crucial role in suppressing inflammatory immune cells and protecting tissues from damage [16, 17, 18], suggesting that IL-10-mediated immune regulation warrants further investigation in the context of C60. For example, the increased IL-10 production could contribute to the generation of regulatory T cells (Tregs) as well as type-I regulatory T cells (Tr1) through the modulation of APC function by C60. IL-10-producing DC, known as DC-10, are well-established as key players in Treg generation [19].

Although our results suggest that the TLR (TLR2, 4, and 9)-MyD88 pathway is a critical mechanism for DC activation, we have not yet identified the specific substances in C60 that trigger these stimulatory signals in DCs. The bacterial cell is composed of various organic molecules with distinct biochemical properties, making it challenging to pinpoint the exact component responsible for DC cytokine production [20, 21, 22]. It remains unclear whether a single substance orchestrates all the observed responses in DCs or if multiple substances are involved in modulating the function. Our previous research showed that C60 structural components (cell wall origin) modulated macrophage function in anti-tumor immunity, contributing to reduced tumor growth in mouse melanoma and breast cancer models [23]. However, the isolation methods in that study were not stringent. Future research will focus on identifying the specific substance(s) in C60 responsible for modulating myeloid cell function. If successful, these substances could have potential applications as dietary supplements or future therapeutics.

As a novel function of C60, we discovered that this probiotic strain can enhance IgA production by stimulating intestinal B cells through DC-derived cytokines. This finding diverges from our earlier works, which focused on C60’s role in activating effector T cells via increased cytokine production and antigen presentation [6, 7, 8]. We conclude that the C60-mediatedly increased IgA production in intestinal B cells is primarily driven by elevated IL-6 and IL-10 productions from stimulated DCs. In vitro culture of PP-derived cells revealed that neutralizing IL-6 and/or IL-10 significantly reduced IgA production, highlighting the crucial role of these cytokines in IgA production. Furthermore, co-culture of PP-derived B cells and DCs demonstrated that the absence of DCs resulted in a failure to produce substantial levels of IgA in HK-C60 stimulation. Our previous studies using human peripheral blood mononuclear cells (PBMCs) also showed that DCs stimulated with *Pediococcus acidilactici* K15 produced IL-6 and IL-10, and these cytokines were essential for IgA production in human B cells [9]. This supports our current findings in the case of C60. Another important observation was that C60 directly promoted B cell proliferation, but IgA production only increased in the presence of DC cytokines. This suggests that IgA upregulation in the intestinal PPs involves at least two distinct steps: B cell proliferation and subsequent exposure to DC cytokines.

To deepen our understanding, additional factors involved in C60-mediated IgA upregulation should be explored beyond IL-6 and IL-10. For instance, B cell-activating factor (BAFF) and a proliferation-inducing ligand (APRIL) from DCs, as well as B cell activation via CD40-CD40 ligand (CD40L) interactions, may influence IgA levels in PPs [24, 25, 26]. Additionally, to examine antigen-specific T cell-dependent IgA production, the activity of T follicular helper (T_FH_) cells should be characterized in C60-administered mice intestinal environment [26, 27]. Since we have already shown that C60 stimulation enhances cytokine production and antigen-presenting activities in APCs, it is plausible that T_FH_ cells may also be affected by these functionally modified myeloid cells. If T cell-dependent IgA production is enhanced by C60, it could lead to the production of high-affinity IgA, contributing to a more targeted defense against pathogenic invasion.

In summary, our studies reveal that C60 is a multipotent probiotic LAB strain with the potential to modulate a wide range of immune responses. While LAB probiotics have yet to reach the therapeutic efficacy needed to replace current medications, further detailed studies may uncover additional, unanticipated benefits, positioning C60 as a candidate for future natural therapeutic applications.

## Acknowledgments

We thank National Agriculture and Food Research Organization (NARO) for kindly providing *Lactococcus lactis* subsp.c*remoris* C60.

## Author Contributions

S.S., A.O., N.K., and T.M. performed experiments. S.S., and N,M,T. established methodologies. S.S., A.O. and N.K. performed data analyses. A.O. and N.M.T. had responsibility in resources. S.S., A.O. and N,M,T. wrote the manuscript. S.S. and N.M.T. reviewed and finalized the manuscript. All authors participated in the discussion. N,M,T. supervised this study.

## Data availability

Corresponding author will provide original data used in this manuscript upon reasonable request.

