## Supplemental Materials for "*Lactococcus lactis* subsp. *cremoris* C60 promotes immunoglobulin A production from B cells through functional modification of dendritic cells in intestinal environment"

Noriko M. Tsuji, Ph.D.


---

**Supplemental Table 1. Primer sequences in real-time PCR analysis**

| Target | Forward (5'-3') | Reverse (5'-3') |
|---|---|---|
| *Il1b* | GCAACTGTTCCTGAACTCAACT | ATCTTTTGGGGTCCGTCAACT |
| *Il4* | GGTCTCAACCCCCAGCTAGT | GCCGATGATCTCTCTCAAGTGAT |
| *Il6* | CCAAGAGGTGAGTGCTTCCC | CTGTTGTTCAGACTCTCTCCCT |
| *Il8* | CAAGGCTGGTCCATGCTCC | TGCTATCACTTCCTTTCTGTTGC |
| *Il10* | GCTCTTACTGACTGGCATGAG | CGCAGCTCTAGGAGCATGTG |
| *Il12a* | CTGTGCCTTGGTAGCATCTATG | GCAGAGTCTCGCCATTATGATTC |
| *Il12b* | TGGTTTGCCATCGTTTTGCTG | ACAGGTGAGGTTCACTGTTTCT |
| *Il23a* | ATGCTGGATTGCAGAGCAGTA | ACGGGGCACATTATTTTTAGTCT |
| *Ifna* | GGATGTGACCTTCCTCAGACTC | ACCTTCTCCTGCGGGAATCCAA |
| *Ifnb* | CAGCTCCAAGAAAGGACGAAC | GGCAGTGTAACTCTTCTGCAT |
| *Ifng* | ATGAACGCTACACACTGCATC | CCATCCTTTTGCCAGTTCCTC |
| *Tnfa* | GACGTGGAACTGGCAGAAGAG | TTGGTGGTTTGTGAGTGTGAG |
| *Tgfb* | CTCCCGTGGCTTCTAGTGC | GCCTTAGTTTGGACAGGATCTG |
| *Tlr1* | TGAGGGTCCTGATAATGTCCTAC | AGAGGTCCAAATGCTTGAGGC |
| *Tlr2* | GCAAACGCTGTTCTGCTCAG | AGGCGTCTCCCTCTATTGTATT |
| *Tlr3* | GTGAGATACAACGTAGCTGACTG | TCCTGCATCCAAGATAGCAAGT |
| *Tlr4* | ATGGCATGGCTTACACCACC | GAGGCCAATTTTGTCTCCACA |
| *Tlr5* | GCCCCGTGTTGGTAATATCTC | ATCTGGGTGAGGTTACAGCCT |
| *Tlr6* | TGAGCCAAGACAGAAAACCCA | GGGACATGAGTAAGGTTCCTGTT |
| *Tlr7* | ATGTGGACACGGAAGAGACAA | GGTAAGGGTAAGATTGGTGGTG |
| *Tlr8* | GAAAACATGCCCCCTCAGTCA | CGTCACAAGGATAGCTTCTGGAA |
| *Tlr9* | ATGGTTCTCCGTCGAAGGACT | GAGGCTTCAGCTCACAGGG |
| *Gapdh* | TGTGTCCGTCGTGGATCTGA | TTGCTGTTGAAGTCGCAGGAG |

### Supplemental Figures

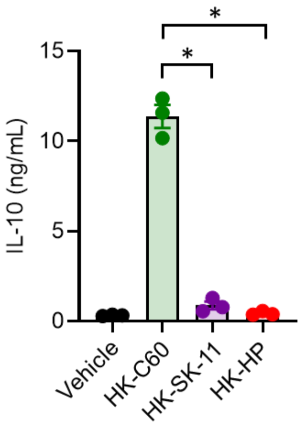


**Supplemental Figure 1.** ***In vitro*** **IL-10 production assay in BMDCs**

BMDCs ($1.0x10^6$/mL) were cultured with vehicle (PBS), HK-C60 ($5.0x10^8$ CFU/mL), HK-SK-11 ($5.0x10^8$ CFU/mL) or HK-HP ($5.0x10^8$ CFU/mL) at 37°C for 24 h. The IL-10 concentration in cultured medium was measured by ELISA. The data were shown as mean ± SEM of three samples from independent experiments. One-way ANOVA was used to analyze data for significant differences. Values of *$p$ < 0.001 were regarded as significant.

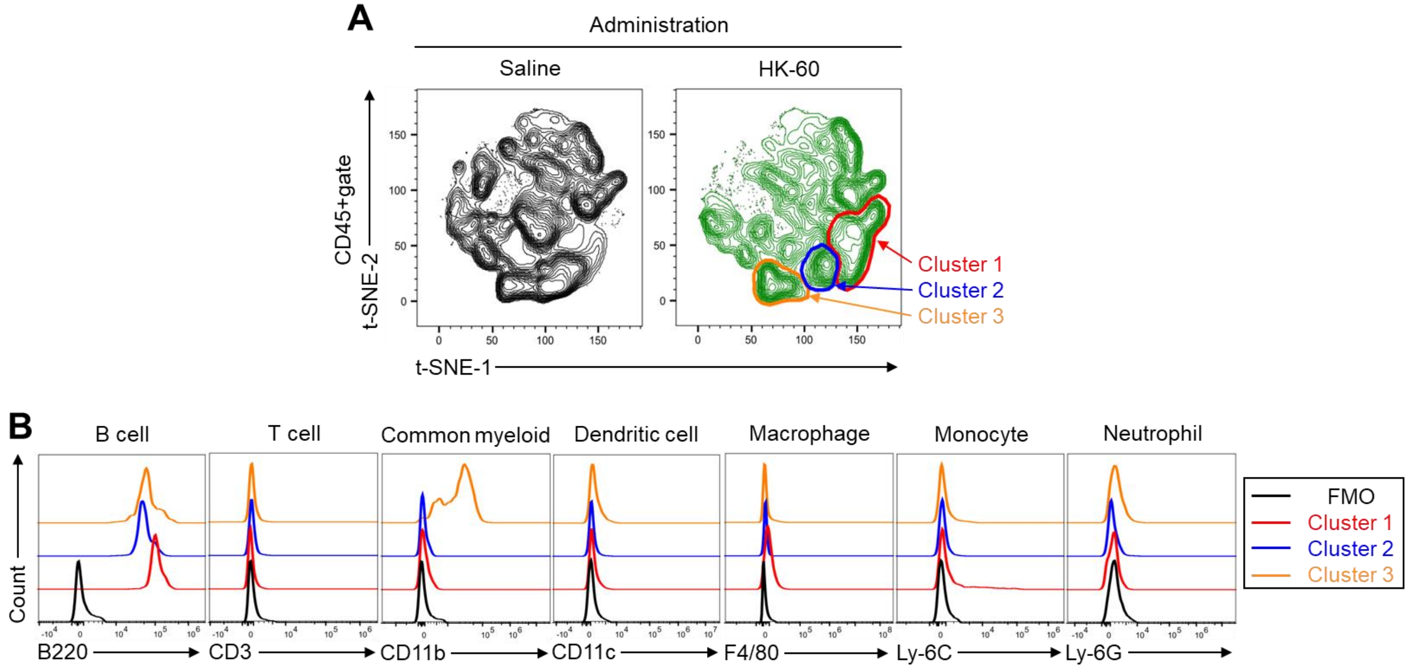


**Supplemental Figure 2. Global immunophenotyping in small intestinal PP**

The mice received i.g. administration of saline or HK-C60 for 2 weeks (n=3 in each group). The small intestinal PP-derived cells were subjected to flow cytometry analysis. A) A t-SNE plot generated by PP-derived cell staining multi markers targeting immune cells. Total three clusters were identified as unique populations in HK-C60 administered group. B) Independent identification of cell type following specific marker expression in three unique clusters.

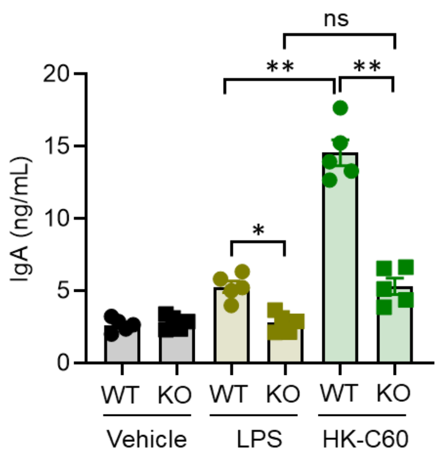


**Supplemental Figure 3. *In vitro* IgA production assay in PP-derived cells originated from MyD88-KO mice**

Small intestinal PP-derived cells were prepared from naïve WT mice or MyD88-KO mice (KO) (n=5 in each group), and the cells ($1.0x10^7$/mL) were subjected to *in vitro* culture with vehicle (PBS), LPS (1 µg/mL) or HK-C60 ($5.0x10^8$ CFU/mL) at 37°C for 72 h. The IgA concentration in cultured medium was measured by ELISA. The data were shown as mean ± SEM of five samples from independent experiments. One-way ANOVA was used to analyze data for significant differences. Values of *$p < 0.01$ and $p$**< 0.001 were regarded as significant. ns: not significant.

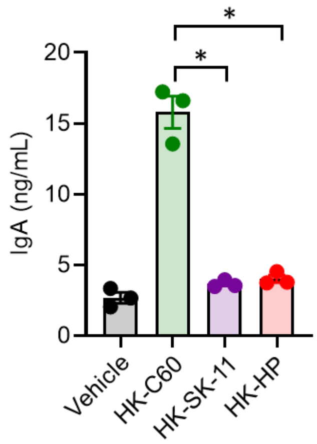


**Supplemental Figure 4. *In vitro* IgA production assay in PP-derived cells**

Small intestinal PP-derived cells were prepared from naïve WT mice (n=3), and the cells ($1.0x10^7$/mL) were subjected to *in vitro* culture with vehicle (PBS), HK-C60 ($5.0x10^8$ CFU/mL), HK-SK-11 ($5.0x10^8$ CFU/mL) or HK-HP ($5.0x10^8$ CFU/mL) at 37°C for 72 h. The IgA concentration in cultured medium was measured by ELISA. The data were shown as mean ± SEM of three samples from independent experiments. One-way ANOVA was used to analyze data for significant differences. Values of *$p <$ 0.001 were regarded as significant.
